# Vector diversity and malaria prevalence: global trends and local determinants

**DOI:** 10.1101/2022.10.13.512182

**Authors:** Amber Gigi Hoi, Benjamin Gilbert, Nicole Mideo

**Author notes:** Corresponding author, Address: 600 Peter Morand Crescent, Ottawa, Ontario, Canada, K1G 5Z3.

## Abstract

Identifying determinants of global infectious disease burden is a central goal of disease ecology. While it is widely accepted that host diversity structures parasite diversity and prevalence across large spatial scales, the influence of vector diversity on disease risk has rarely been examined despite the role of vectors as obligatory intermediate hosts for many parasites. Malaria, for instance, can be transmitted by over 70 species of mosquitoes, but the impact of this diversity on malaria risk remains unclear. Further, such relationships are likely dependent on the context in which disease transmission occurs, as arthropod life history and behavior are highly sensitive to environmental factors such as temperature. We studied the relationship between vector diversity, malaria prevalence, and environmental attributes using a unique dataset we curated by integrating several open-access sources. Globally, the association between vector species richness and malaria prevalence differed by latitude, indicating that this relationship is strongly dependent on underlying environmental conditions. Structural equation models further revealed different processes by which the environment impacts vector community assemblage and function, and subsequently disease prevalence, in different regions. In Africa, the environment exerted a top-down influence on disease through its role in shaping vector communities, whereas in Southeast Asia, disease prevalence is influenced by more complex interactions between the physical and socioeconomic environment (i.e., rainfall and GDP) and vector diversity across sites. This work highlights the key role of vector diversity in structuring disease distribution at large spatial scales and offers crucial insights to vector management and disease control.

**Significance statement:** The global health threat from persistent and emerging vector-borne diseases continues to increase and is exacerbated by rapid environmental and societal change. Predicting how disease burden will shift in response to these changes necessitates a clear understanding of existing determinants of disease risk. We focused on an underappreciated potential source of variation in disease burden – vector diversity – and its role in structuring global malaria distribution. Our work revealed that vector diversity influences malaria prevalence and that the strength and nature of this association strongly depend on local environmental context. Extending disease transmission theory, surveillance, and control to embrace heterogeneity in vector community structure and function across space and time is an asset in the fight against vector-borne diseases.

## Introduction

Global infectious disease burden remains high despite decades of control efforts (Lozano et al. 2012). New diseases are emerging at a rapid pace, and many existing diseases persist (Smith et al. 2014; Bloom and Cadarette 2019). These threats to public and global health are not distributed evenly around the world but tend to be concentrated in the tropics (Guernier et al. 2004; Bonds et al. 2012). Specifically, there is a distinct latitudinal gradient of increasing infectious disease burden from the poles towards the equator, both in terms of disease prevalence (Bonds et al. 2012) and the numbers of distinct diseases (Guernier et al. 2004; Dunn et al. 2010; but see Kamiya et al. 2014).

Latitude has long served as a powerful proxy for capturing the myriad factors that covary along these clines and shape large-scale biological patterns (Pianka 1966; Hillebrand 2004), including disease distributions (Stephens et al. 2016; Murray et al. 2018). Variation in biotic (host abundance and diversity), abiotic (climate and topography), and anthropogenic (socioeconomics) factors across latitude is thought to contribute to the observed gradients in disease diversity and prevalence (Murray et al. 2018). These factors may interact and influence one another in a complex manner, making it difficult to determine their relative importance (Dunn et al. 2010; Franklinos et al. 2019). Further, latitude as a proxy obscures much of the heterogeneity in the eco-epidemiological context that exists along each cline. For instance, Indonesia and Kenya both lie on the equator, but share few other features: in terms of climate, Indonesia is mostly situated in the tropical rain forest biome, where temperature and rainfall are consistently high year round, whereas Kenya spans seasonal rain forest and shrubland biomes, characterized by distinct wet and dry seasons (Kaplan et al. 2003). Moving beyond latitude to elucidate the general ecological principles that contribute to patterns of disease across diverse settings is a central goal in disease ecology and epidemiology, and is crucial to informing current disease control policy and predicting future outbreaks.

The intimate relationship between parasites and their hosts lies at the core of the network of interactions that give rise to global disease patterns (Poulin 2014). Parasite diversity and prevalence is (in part) a direct consequence of host diversity and prevalence, as by definition parasites require their hosts for survival and dispersal (Guernier et al. 2004; Dunn et al. 2010). Indeed, global analyses suggest parasite diversity tracks well-established latitudinal gradients in mammalian and avian diversity for which species richness is highest at the equator and decreases towards the poles (Guernier et al. 2004; Dunn et al. 2010; Murray et al. 2015). Host diversity is in turn thought to be structured by the abiotic environment: the tropics consist of environments conducive to supporting large and diverse host communities through high energy and water supply, coupled with high environmental heterogeneity (“climatically-based energy hypothesis”; Guernier et al., 2004). Anthropogenic factors such as socioeconomic development can have multifaceted effects on infectious disease outcome and risk: investment into health care can improve disease outcome directly, whereas land-use change due to economic activity may lead to changes in local host community assembly (e.g., altered habitat favoring a particular host species), indirectly affecting disease risk (Lambin et al. 2010; Gottdenker et al. 2014; Wood et al. 2017; Gibb et al. 2020). The effects of the natural and man-made environments on disease are thus frequently formulated as acting through hosts in a top-down, hierarchical manner (Fig. 1A, top; Cumming & Guégan, 2006). Alternatively, the environment may induce changes in host or vector behavior and life history, and subsequently exposure to disease, independent of any effect on local community structure (Fig 1A, bottom). For instance, on average, humans engage the outdoors more often when it is warm and sunny, increasing exposure to disease vectors such as ticks and mosquitoes (e.g., Lambin et al., 2010; Wagner et al., 2020). Economic incentives may also drive host movement in and out of endemic regions on short timescales (e.g., migrant workers), sparking outbreaks and contributing to seasonal disease cycles (Moyce and Schenker 2018).

**Fig 1.**
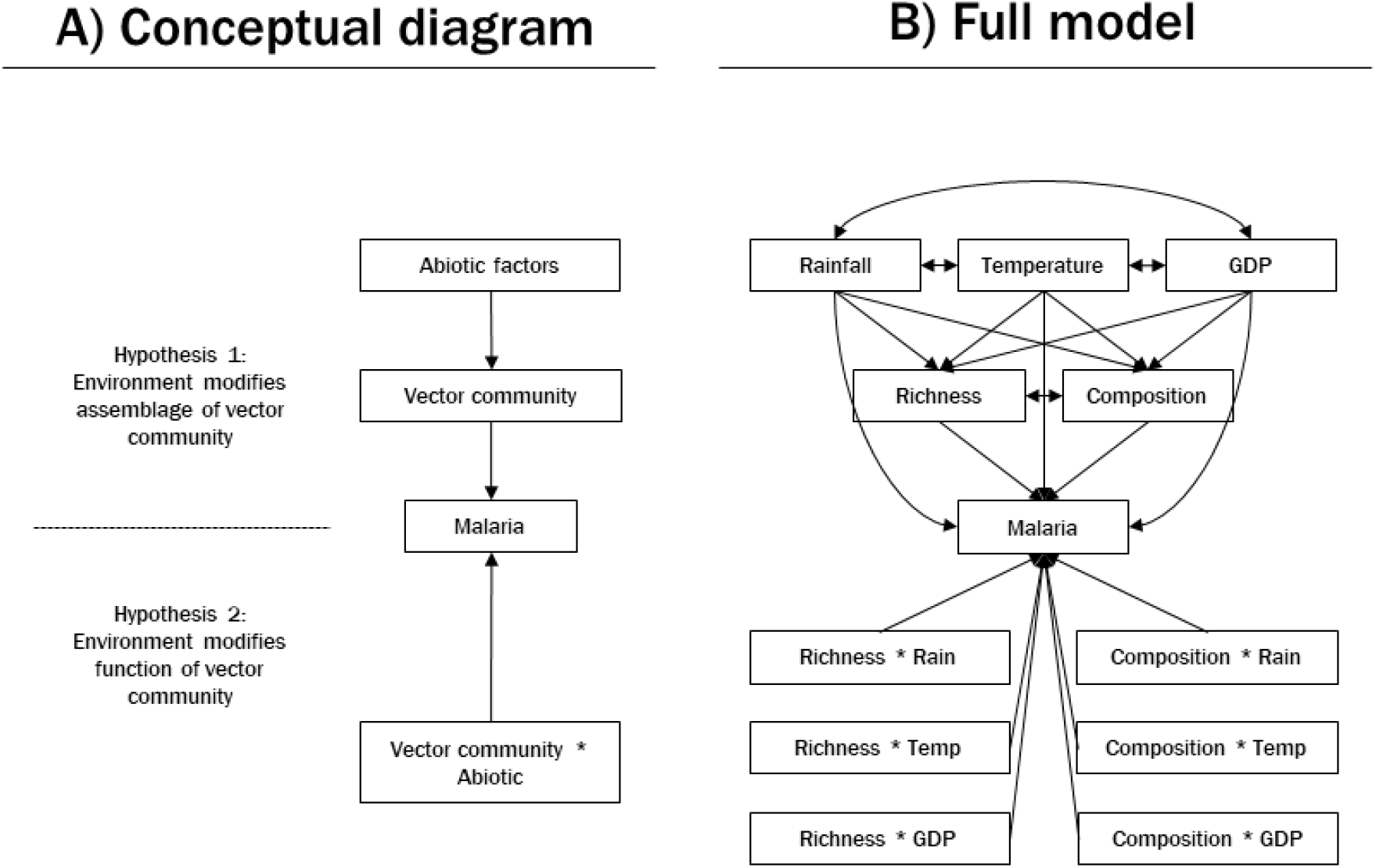
Path diagram representing hypothesized relationships between characteristics of the abiotic environment, vector community attributes, and malaria prevalence. A) The environment may influence disease risk via vector community assemblage (Hypothesis 1) or function (Hypothesis 2). B) Path diagram for hypothesized full model. Paths (i.e., arrows) represent the direction of the hypothesized causal relationship between variables, with the arrow pointing from the predictor to the response. Double-headed arrows indicate hypothesized correlation between variables.

The impacts of environment on disease spread are particularly prominent in vector-borne disease systems (Guernier et al. 2004). Arthropod vectors are often obligatory intermediate hosts of parasites, and it is widely accepted that vector traits such as biting and mortality rates play disproportionate roles in driving disease dynamics (Koella 1999). These traits are under strong regulation by the environment since, as exotherms, the vital and activity rates of arthropods are temperature dependent (Shapiro et al. 2017; Mordecai et al. 2019; Cator et al. 2020). Temperature regulated life-history traits also determine vector competency (Paaijmans et al. 2012, 2013; Brady et al. 2016; Ferraguti et al. 2018), and scale up to influence vector population dynamics (Beck-Johnson et al. 2013, 2017) and species distribution (Sinka et al., 2010; Sinka et al., 2011). Many mathematical models of vector-borne disease transmission therefore leverage temperature reaction norms to predict vector abundance and disease risk, and have been applied to systems including Malaria (e.g., Mordecai et al., 2013), Dengue (e.g., Kamiya et al., 2020), and Zika (e.g., Ryan et al., 2020) (see Mordecai et al., 2019 for a recent review of such models).

While vector abundance is an important determinant of disease spread, many parasites can be transmitted by multiple vector species, and these vector communities are rarely homogeneous across space and time. Human malaria parasites, for example, predominantly infect just one vertebrate host in natural systems (occasionally also infecting other primates; Faust & Dobson, 2015) but can utilize more than 70 species of mosquitoes as vectors (Hay et al. 2010). Comparing mosquito communities around the globe, the precise number of species and their relative abundances vary considerably (e.g., Kulkarni et al., 2006; Oo et al., 2003). These mosquito species also encompass substantial diversity in seasonal activity, habitat, and feeding preferences (Massey et al., 2016; Sinka et al., 2010, 2011), such that different assemblages of vectors should have differential impacts on transmission dynamics (Hoi et al. 2020). Local analyses suggest vector diversity amplifies malaria prevalence (Hoi et al. 2020), but it is unclear whether this association is generalizable over large spatial scales and various environmental contexts under which vector-host interactions occur (Stephens et al. 2016; Murray et al. 2018). At the macroecological scale, mosquito diversity conforms to a latitudinal diversity gradient (Hillebrand 2004; Foley et al. 2007) parallel to that of malaria burden (Gallup and Sachs 2001), where each increases in magnitude towards the equator. However, as far as we are aware, latitudinal gradients have not been investigated for the subset of mosquito species that are competent vectors – a distinction that is important for predicting how vector diversity should influence disease risk. Further, while habitat suitability is fundamental to the presence of individual vector species (e.g., Hay et al., 2010), it is unclear if local environmental filters may also promote different mosquito vector community assemblages (Fig 1A, top; Ferraguti et al., 2016) or modify the disease transmission capacity of those communities (Fig 1A, bottom; Park et al., 2015). These two pathways have been demonstrated independently in tick-borne parasites of primates in Africa (Cumming and Guégan 2006) and midge-borne viruses of wild ungulates in south-eastern United States (Park et al. 2015), however, few studies have examined them in tandem and investigated their relative contribution to shaping disease patterns.

A clear vision of how biotic and abiotic drivers of disease intersect is an asset in combating infectious diseases in the ever-changing Anthropocene (Harvell et al. 2002; Franklinos et al. 2019). This study seeks to elucidate the complex factors that underpin global patterns of malaria burden, with an explicit focus on natural patterns of vector diversity. Our goal is to bolster our current understanding of environmental effects on vector abundance and disease (e.g., Mordecai et al. 2019) to consider the role of the broader diversity of vectors that transmit malaria and different pathways by which the environment may modify entomological risk. We build our analysis in multiple stages to capture both macroecological trends and general patterns in local disease drivers. First, we investigate whether the global occurrence of mosquito vector species is consistent with expectations of a latitudinal gradient. Next, we determine whether vector species richness is associated with malaria prevalence, and whether this association is homogenous across latitudes. Our approach allows us to assess whether global patterns hold at smaller scales or if the relationship between vector diversity and malaria prevalence is context dependent. Finally, we adopt a structural equation modelling framework to elucidate pathways through which the environment influences malaria transmission in the major malaria-endemic regions of sub-Saharan Africa and South Asia, Southeast Asia, and Pacific Islands (hereafter Southeast Asia). Malaria is prevalent in sub-Saharan Africa, with young children suffering mild to severe symptoms while infections in adults are often asymptomatic (i.e., holoendemic), whereas in Southeast Asia, malaria transmission is less stable and largely driven by seasonal outbreaks (i.e., hypoendemic) (Baird, 2017; Carter & Mendis, 2002; Table 1; Fig. A1 in Appendix A). These regions also broadly differ in climate and landscape, as well as the biology of the hosts, vectors, and parasites present (Baird, 2017; Carter & Mendis, 2002; Table 1; Fig. A1 in Appendix A). In light of these distinctions, we assess the influence of the environment on vector communities and malaria independently in each region.

**Table 1.**
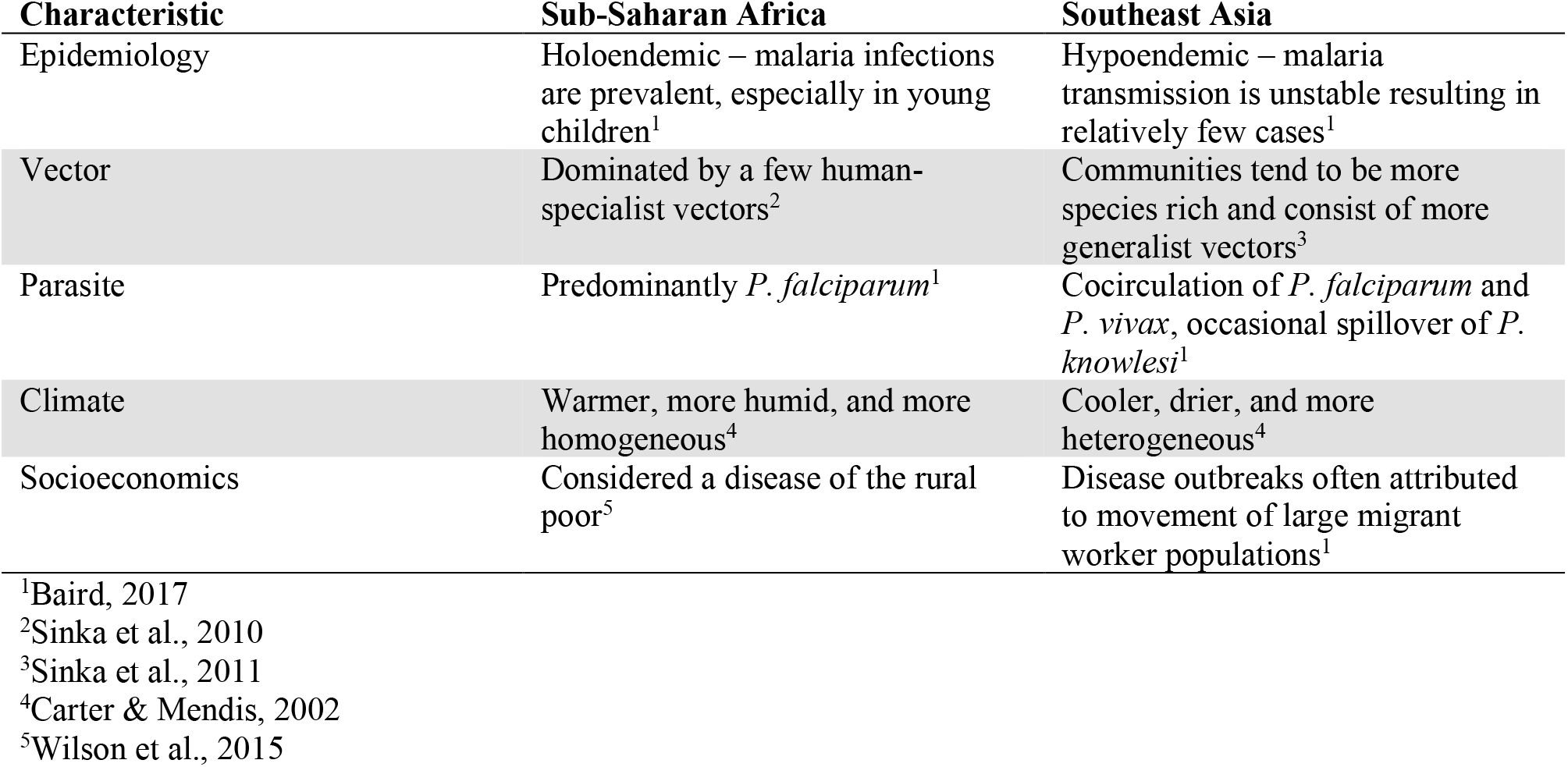
Epidemiological, biological, environmental, and socioeconomic characteristics of the malaria transmission context in sub-Saharan Africa and Southeast Asia. In general, there are broad differences in the ecological and socioeconomical background of the two continents, as well as substantial intra-continental variation in these factors. We focused on the former with the aim of providing a foundation for our analysis plan and for the interpretation of our results.

## Results

### Overview of data

Our curated dataset consisted of information on malaria prevalence, vector communities, and study site attributes integrated from several open-access data sources (for details, see Methods: Data sources and management). We defined malaria as infections caused by any, or combinations, of the five human malaria parasite species (*Plasmodium falciparum, P. vivax, P. malariae, P. ovale*, and *P. knowlesi*). The structure of vector communities were represented by species richness and proportion of all species that are primary vectors, defined as those vector species that contribute significantly to malaria transmission (Hay and Snow 2006; Service 2012). We also included average monthly temperature and rainfall, and sub-national, per capita GDP, as proxies of the physical and social environments, respectively. The final dataset consisted of 133 complete records from 92 locations, spread over 17 countries and 15 years (1990-2005).

Malaria prevalence was highest in sub-Saharan Africa (Fig. 2A). Out of 54 globally recognized primary vectors of malaria, 30 were represented across our study sites. Secondary vectors were found worldwide, and 33 secondary vectors species were added to the dataset. Sub-Saharan Africa, Asia, and South America did not share any vector species. Sites in Asia consisted of more vector species (Fig. 2B) and these communities tended to be made up of a higher proportion of secondary vectors (Fig. 2C). These sites were also slightly cooler, drier, and had higher per capita GDP than sites in Africa. The frequency distributions of environmental variables (temperature, rainfall, GDP), vector community attributes (species richness, composition), and malaria prevalence across sites are presented in Appendix A.

**Fig 2.**
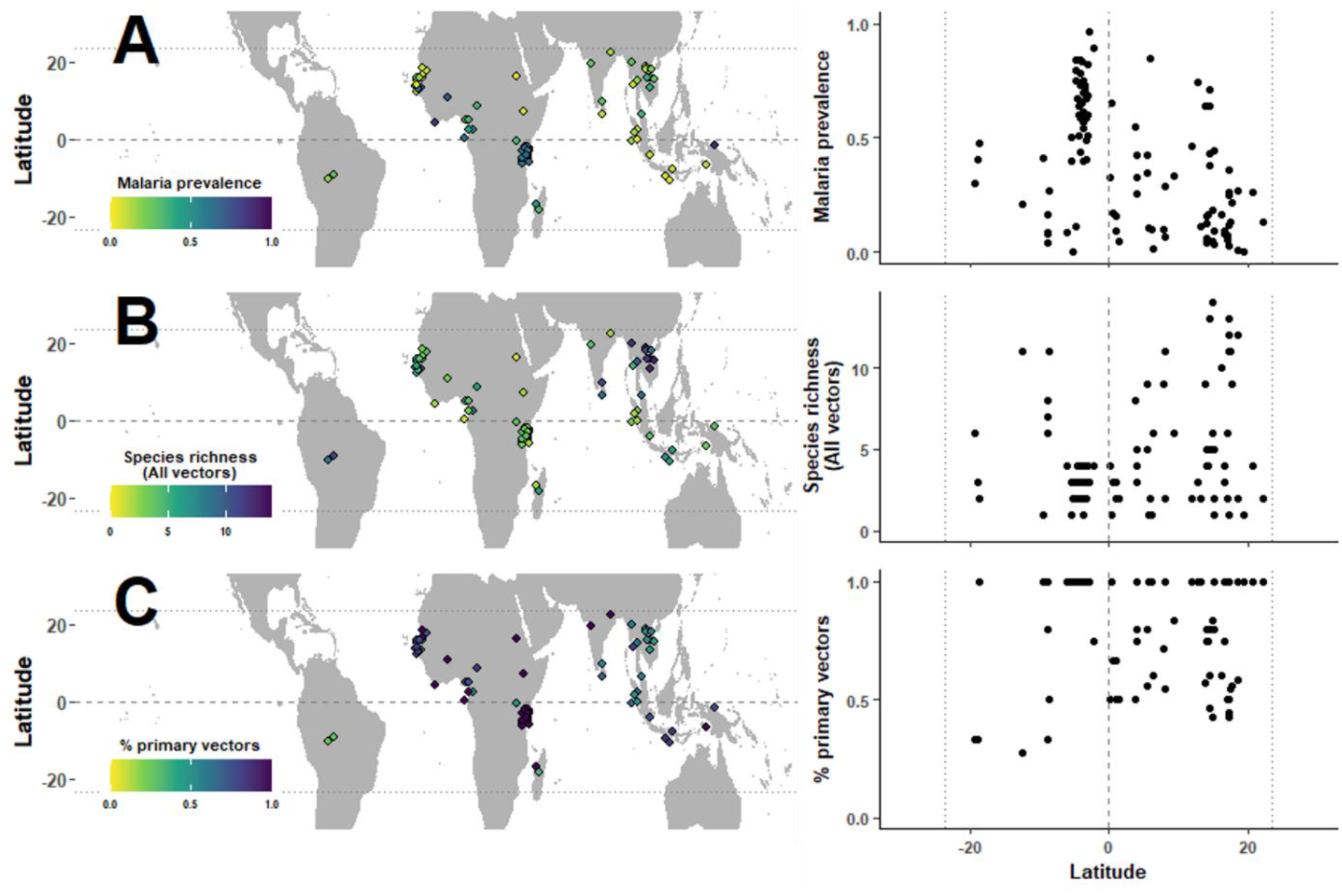
Distribution of A) malaria prevalence, B) total vector species richness, and C) percent primary vectors by species richness across the tropics, 1990-2005. Each data point on maps and scatterplots represents one site (n=92). For sites that were sampled repeatedly over time, the average malaria prevalence and total number of unique vector species observed over the study period was presented. Dashed line represents the equator, and dotted line marks the tropical region (latitude range 23.5°S to 23.5°N) in all plots.

### Global trends in vector diversity and malaria prevalence

Malaria prevalence was negatively correlated with distance from the equator (Pearson’s correlation=-0.43, p<0.01; Fig. 2A), however, vector diversity did not conform to this predicted latitudinal gradient. Vector species richness was positively associated with distance from equator (Pearson’s correlation= 0.49, p<0.01; Fig. 2B), though there appeared to be more primary vectors in the northern hemisphere (Fig. 2C), suggesting that vector diversity in the southern sites was driven by the presence of secondary vectors.

Globally, primary vectors tend to be over-represented in abundance relative to their species richness (Fig. 3). Thirty-four vector communities (located predominantly in east Africa) consisted entirely of primary vectors, and most of the other communities aggregated above the 1:1 line in Fig. 3, indicating that their species compositions were not even, but rather dominated by primary vectors. One interpretation of this pattern is that vector competency is positively correlated with competitive ability. Alternatively, it may simply reflect the fact that being numerically dominant is a key criteria used in the incrimination of primary vectors, regardless of intrinsic competence (Hay et al. 2010). The main exceptions to this observed pattern were several sites in Laos (average distance between sites=62.6 km) in which primary vectors existed in relatively low abundances despite making up ∼50% of vector species.

**Fig 3.**
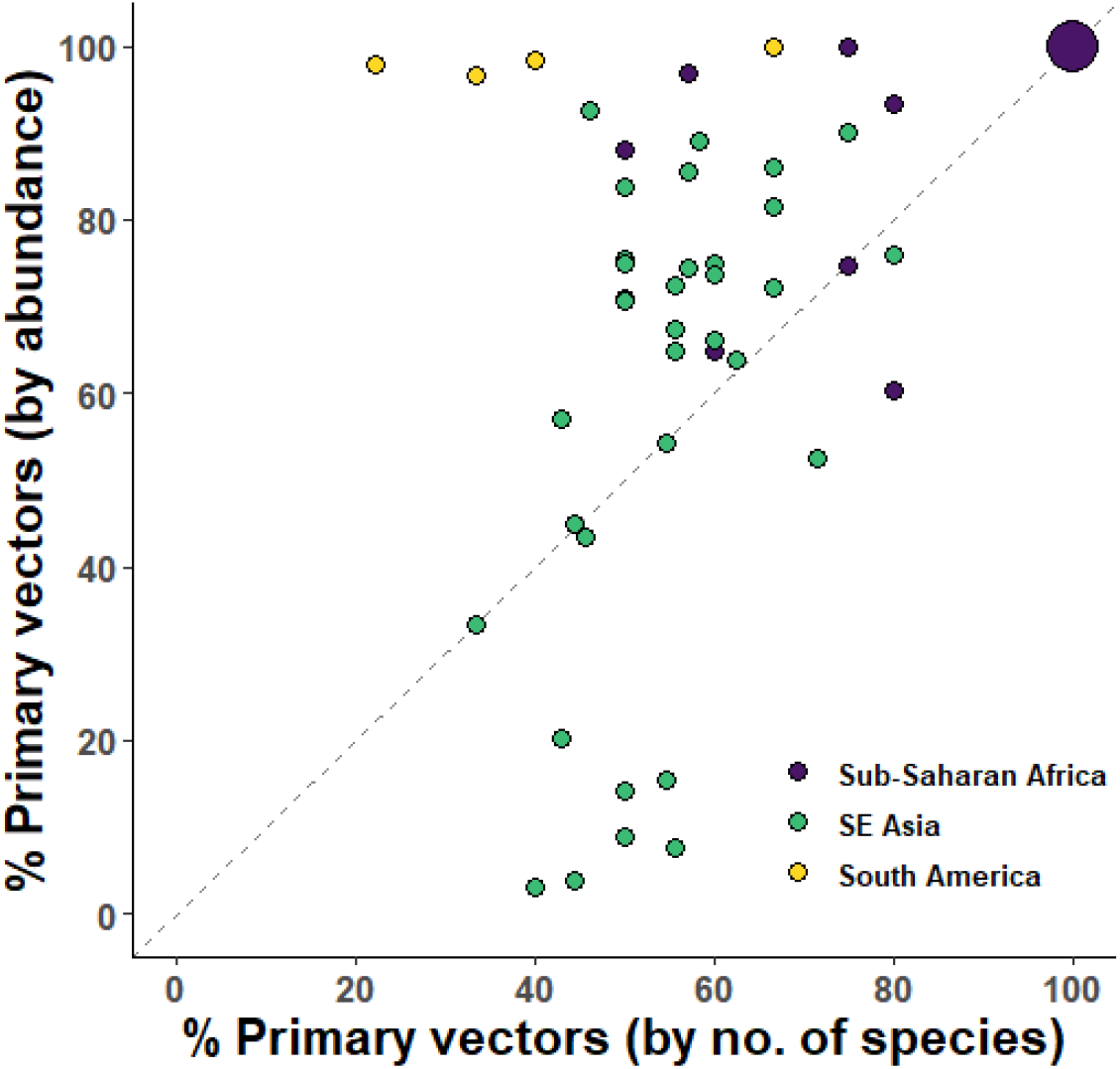
Representation of primary vectors in each community by species richness and abundance relative to all vectors. Each data point represents one observation at the site level and includes a subset of the data (n=83) where species abundance was available, except for the large circle on the top-right corner which represents 34 vector communities in sub-Saharan Africa composed entirely of primary vectors. Dashed line represents a one-to-one relationship between the relative richness and relative abundance of primary vectors. Sites were separated into regions based on continent.

### Latitudinal gradient in vector diversity-disease relationships

We used generalized linear mixed models (GLMMs) to examine the relationship between vector diversity and malaria prevalence at study sites located within the tropics (n=126). This relationship was strongly dependent on latitude, a proxy for environmental conditions known to covary along this gradient (Table 2, Fig 4). The apparent negative association between vector species richness (both primary and total species richness) and malaria prevalence was not statistically significant (Table 2, Fig 4A), however, vector species richness interacted with latitude (absolute values) to influence malaria prevalence (Table 2, Fig 4B). Specifically, there was a strong positive association between vector species richness and malaria prevalence near the equator, but this relationship weakens and reverses moving towards the edge of the tropics (Fig 4B). These effects were weak (fixed effects explained approximately 1% of variation in the data for both models; Table 2) but were captured despite the vast variation between countries and sites and over time (fixed and random effects together explained more than 30% of the variation in the data for both models; Table 2).

**Table 2.**
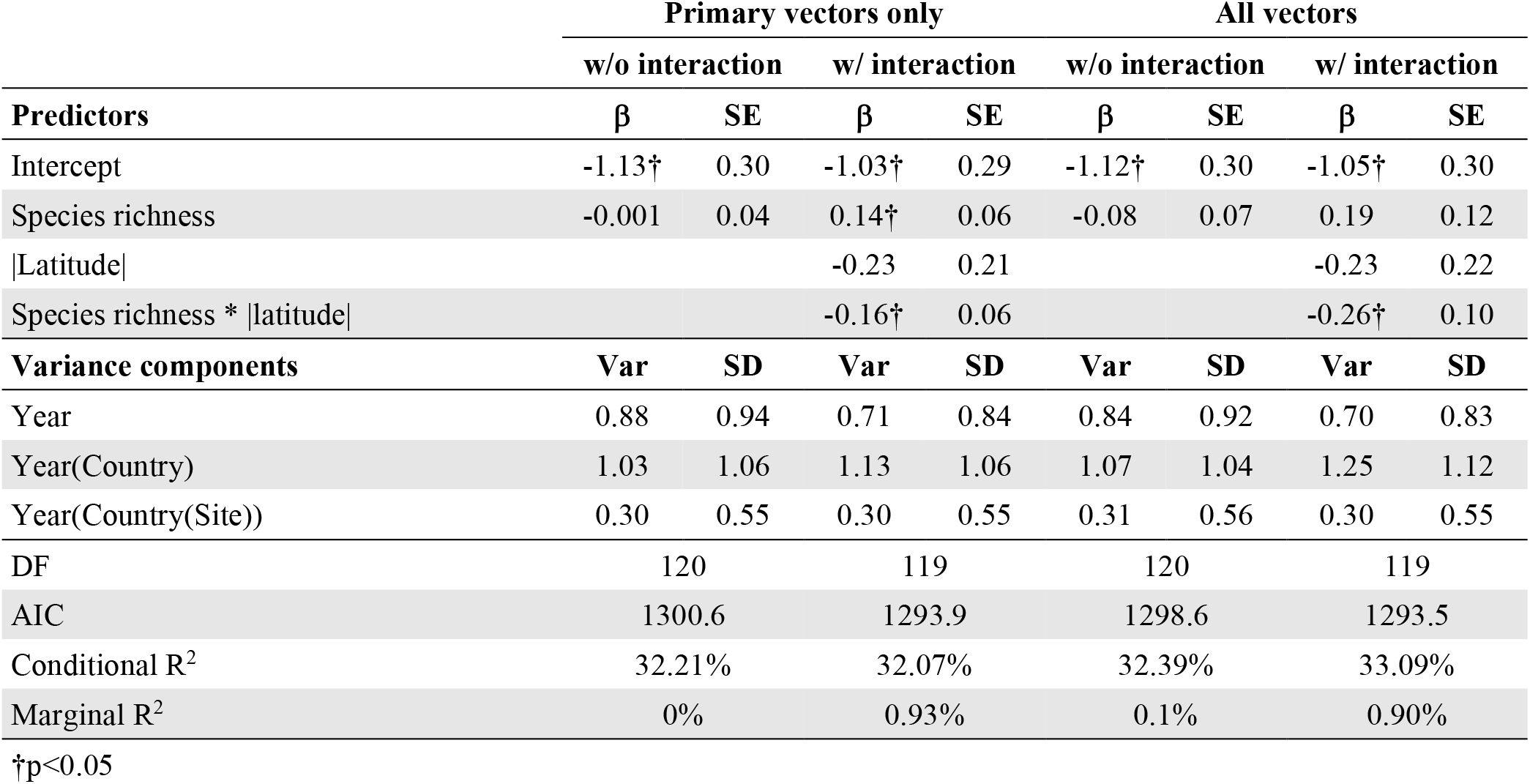
Generalized linear mixed models for malaria prevalence at 126 sites across the tropics. The response variable for all models was malaria prevalence, coded as an aggregate binomial variable and weighted by number of individuals sampled. The following predictor variables were considered: vector species richness (primary vectors only or all vectors), distance from the centroid of each site to the equator (i.e., absolute value of latitude), and their interaction. We constructed a total of four models to investigate the effect of number of primary versus all vectors, and the inclusion versus absence of a species richness by latitude interaction, on malaria prevalence in a factorial manner. All predictors were centered and scaled to unit variance to facilitate model convergence. Several random effects were included to account for non-independence in the data: site (some sites were measured repeatedly), country, and year, nested in that order. β: regression coefficients. SE: standard error. DF: degrees of freedom. AIC: Akaike information criterion. Conditional R^2^: proportion of variation explained by fixed and random effects combined. Marginal R^2^: proportion of variation explained by fixed effects only. R^2^ statistics were estimated using methods outlined in Nakagawa and Schielzeth 2013, and Nakagawa et al. (2017).

**Fig 4.**
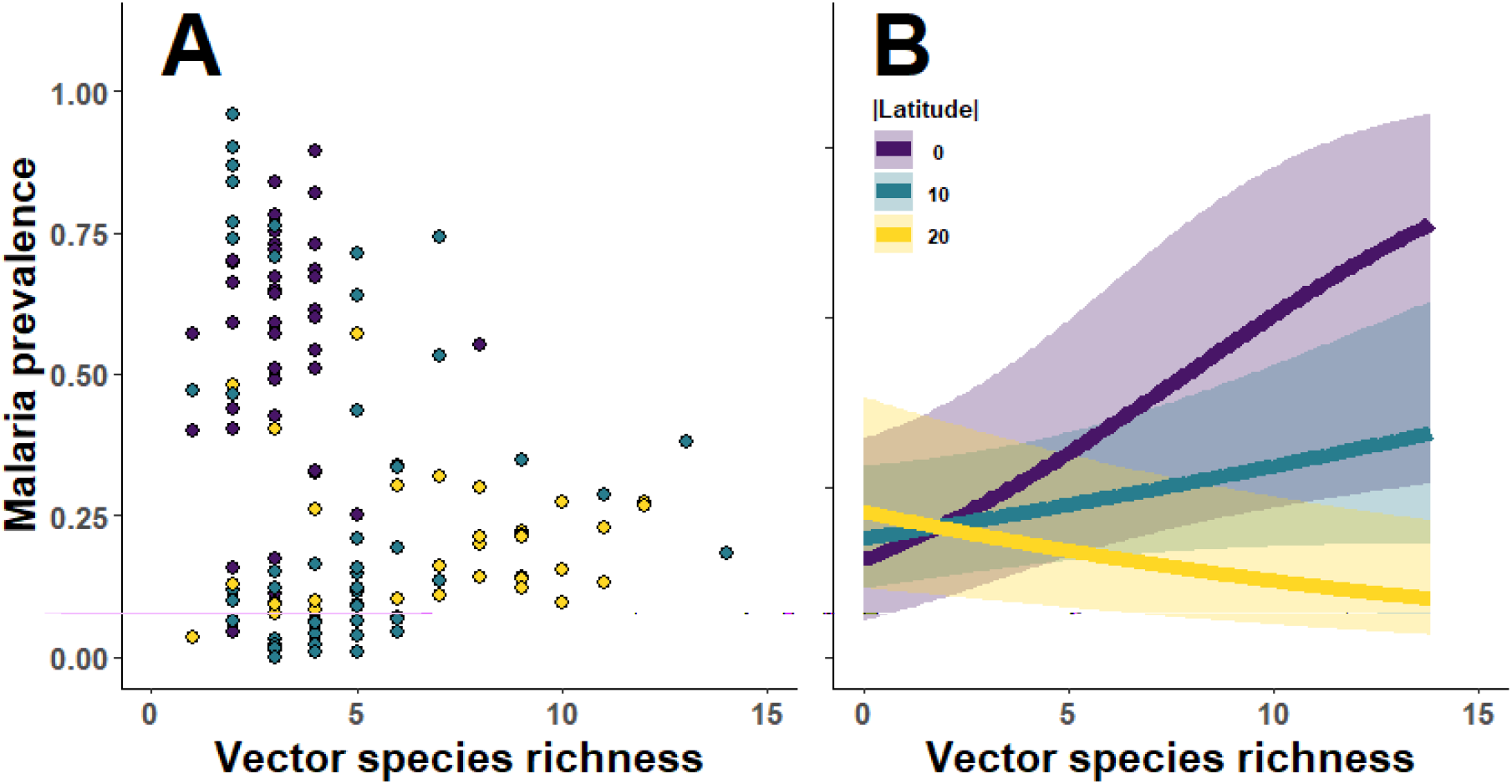
The relationship between malaria prevalence and total vector species richness in the tropics. **A) Scatterplot showing the relationship between malaria prevalence and vector species richness.** Each data point represents one observation (n=126), and the color corresponds to the latitudinal position of the site rounded to the nearest 10 degrees. This association between malaria prevalence and vector species richness was assessed using a generalized linear mixed model and was not statistically significant. **B) Predicted effect of number of vectors on malaria prevalence by distance from equator (absolute latitude)**. Malaria prevalence was modelled using generalized linear mixed models with binomial distribution weighted by the number of individuals sampled. Predictor variables considered were vector species richness, latitudinal position of the centroid of each site, and their interaction. Bands represent 95% confidence interval of prevalence estimates. Both models include study site, country, and year, nested in succession, as a random effect to account for the hierarchical structure of the data. Details of model structure and results for both can be found in Table 2 and the main text.

### Structural equation models revealed regional differences in drivers of malaria burden

We developed a structural equation model (SEM) for an in-depth analysis of environmental variables that mediate the vector diversity-disease relationship at the site level. Broadly, this SEM was designed to distinguish between two hypothesized pathways by which the environment could influence disease risk: 1) exerting a top-down influence on malaria via its effect on vector community assemblage or 2) modifying the disease transmission efficiency of vector communities (Fig 1A). We identified SEMs for the regions of sub-Saharan Africa and Southeast Asia optimized from one *a priori* full model (Fig. 1B) via a predetermined model selection procedure (see Methods: Statistical analysis). The Africa sub-dataset consisted of 64 observations and the Asia sub-dataset included 57. Study sites span the entire latitude range of the tropics in both regions (Fig. 2). Directed separation tests confirmed no important paths missing from either model (Table 3).

**Table 3.**
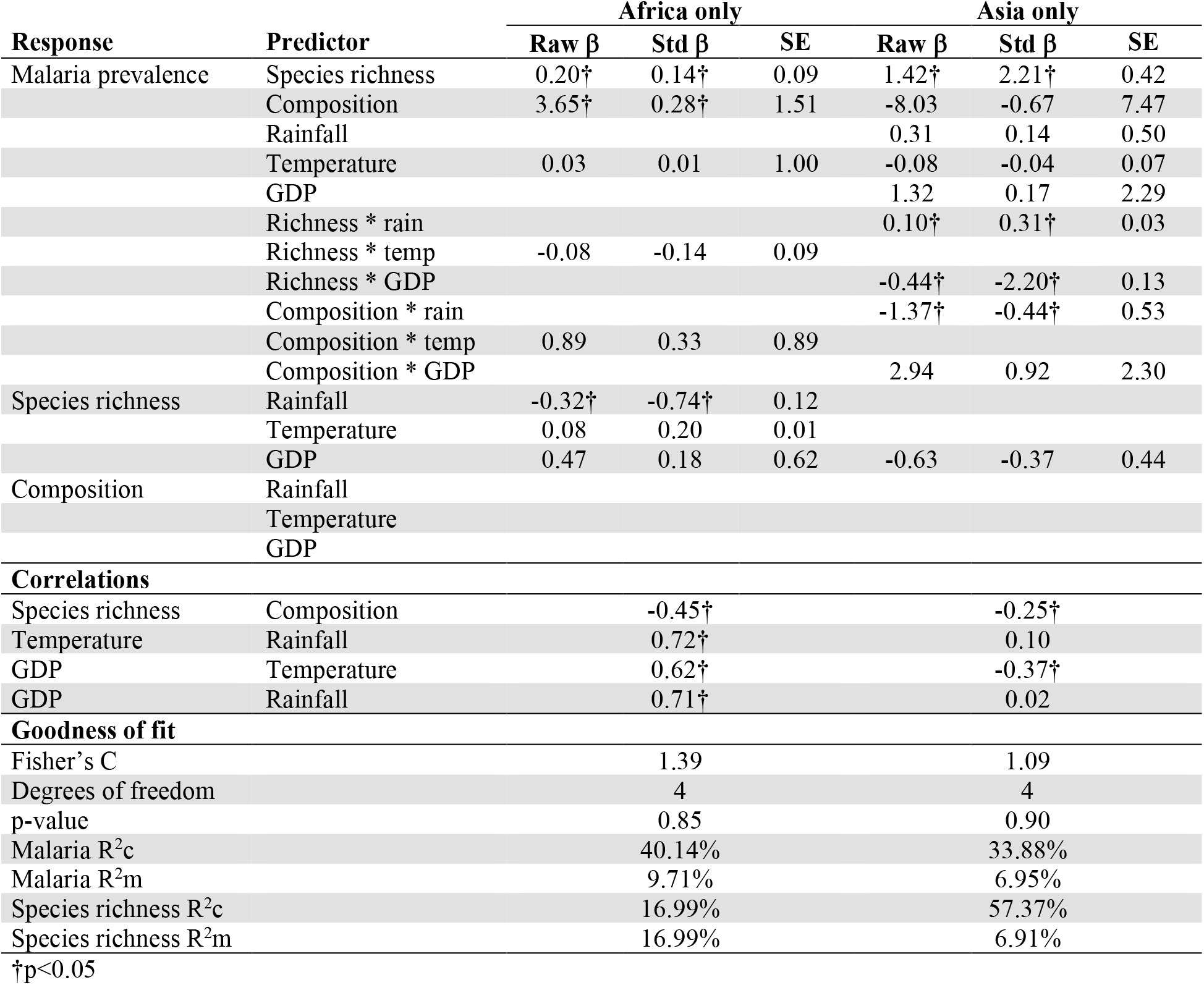
Results for structural equation models fitted to regional (sub-Saharan Africa and Southeast Asia) subsets following model selection. All response variables, along with all corresponding predictors as depicted in Fig. 1B are listed below. All sub-models also include a random effect term to account for non-independence in the data: site (some sites were measured repeatedly), country, and year, nested in that order. A blank cell indicates a path that was dropped during model selection (via procedure detailed in main text). Raw effect estimates (Raw β), standardized effect estimates (Std β), and standard errors (SE) are provided. The d-separation test was used to assess model fit, and the associated Fisher’s C statistic, degrees of freedom, and p-value are reported. The null hypothesis of the d-separation test is that there are no important links missing in this model. Coefficients of determination (R^2^) for the focal response variable (malaria prevalence) are given. Conditional R^2^ (R^2^c): proportion of variation explained by fixed and random effects combined. Marginal R^2^ (R^2^m): proportion of variation explained by fixed effects only. R^2^ statistics were estimated using methods outlined in Nakagawa and Schielzeth 2013, and Nakagawa et al. (2017).

We observed regional differences in the factors shaping vector communities and influencing malaria prevalence. In sub-Saharan Africa, malaria prevalence was associated with both high vector species richness and high proportion of primary vector species (Table 3, Fig. 5A, Fig. B1-ii & B1-iii in Appendix B). Since vector species richness was negatively correlated with the proportion of primary vectors in a community, the overall positive effect of vector diversity thus weakens at high species richness with the introduction of secondary vectors (Table 3, Fig. 5A). Rainfall, temperature, and GDP covaried, but rainfall was the only environmental factor directly associated (negatively) with vector species richness (Table 3, Fig. 5A, Fig B1-i in Appendix B). Taken together, sites with more rain, warmer temperatures, and higher GDP tended to house depauperate mosquito communities consisting mostly of primary vectors. The net effect in these sites is a lower malaria prevalence. Most of the interaction terms between biotic and abiotic factors were dropped during model trimming, and the ones retained in the final model (interactions with temperature) were not significant predictors of malaria. This suggested that in sub-Saharan Africa, the environment influenced malaria prevalence in a top-down manner by way of structuring local vector communities (Fig. 1A, Hypothesis 1).

**Fig 5.**
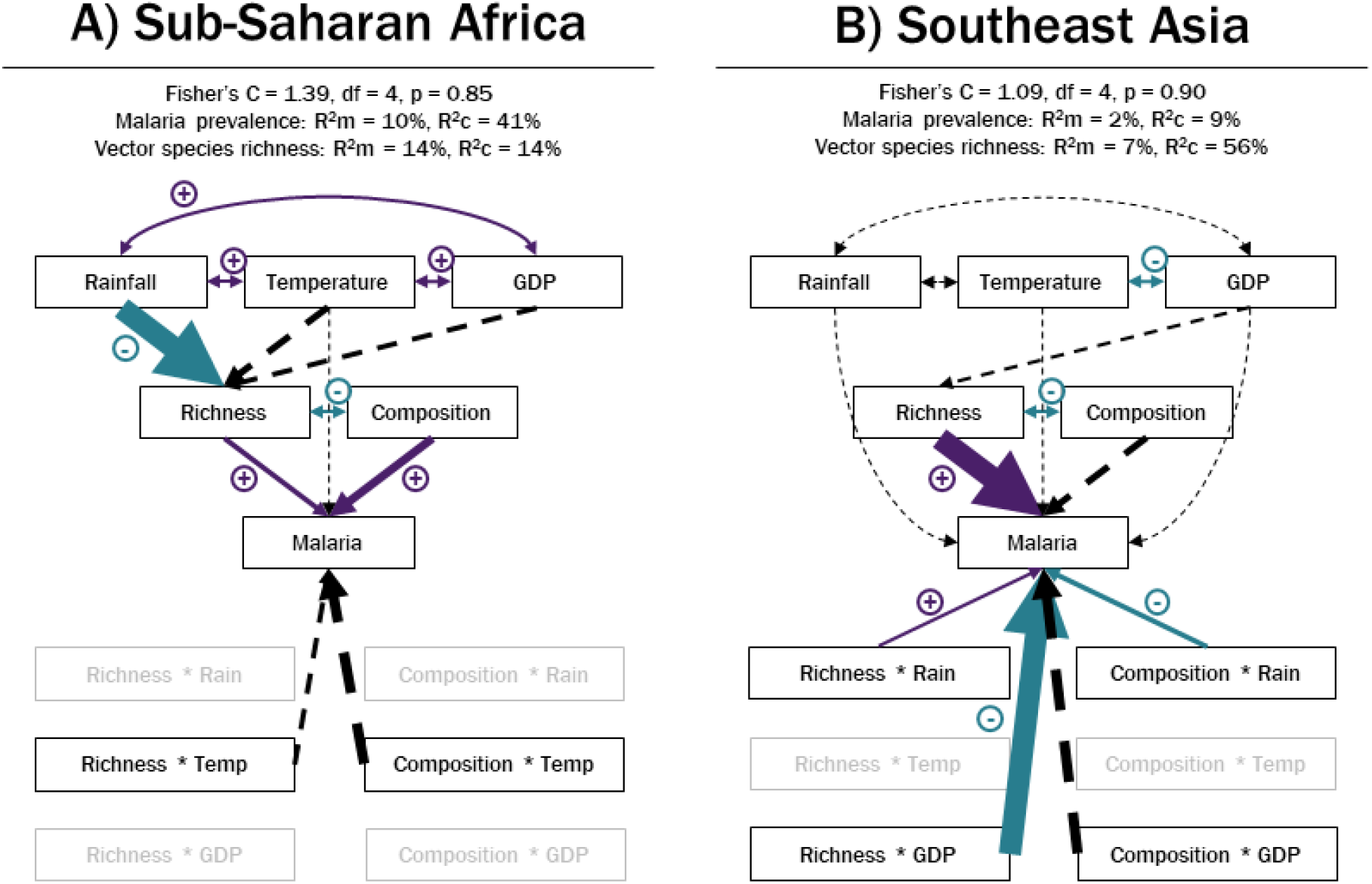
Path diagrams depicting model selection and fit results for A) sub-Saharan Africa (n=64) and B) Southeast Asia (n=57). Model selection procedure detailed in the main text. Paths (i.e., arrows) represent the direction of the modelled causal relationship between variables, with the arrow pointing from the predictor to the response. Arrow size is proportional to the strength of paths within each model but not across models. Double-headed arrows indicate correlations and are not scaled to effect size. Purple arrows represent positive associations, teal represent negative associations, and dashed arrows are associations that were not statistically significant. Variables in grey were dropped during model trimming. See Table 2 for details of model fit results.

Results were drastically different in Southeast Asia. Although species richness still had a significant positive association with malaria prevalence, none of the environmental variables were associated with vector community attributes (Table 3, Fig. 5B). Rather, the environment influenced malaria by altering the effects of vector diversity (vector community × abiotic interactions). Rainfall modified the relationship between vector species richness, composition, and malaria: increasing rainfall tipped a slight negative association between vector species richness and malaria to being positive, but had the opposite effect on the relationship between composition and malaria (Table 3, Fig. 5B, Fig. B2-i and B2-ii in Appendix B). Similarly, the negative interaction between species richness and GDP meant that the positive effect of species richness on malaria weakens and reverses in sites with higher economic output (Table 3, Fig. 5B, Fig. B2-iii in Appendix B). The overall significance of vector community × abiotic interactions in influencing malaria prevalence in Asia suggested that the predominant effect of the environment, especially that of the socioeconomic environment, was to alter the function of vector communities (Fig. 1A, Hypothesis 2).

### Explanatory power of fixed vs. random effect predictors

Random effects maintained high explanatory power across most of our models (Table 2 & 3), indicating that substantial spatial-temporal variation in our modelled response variables was not captured by our fixed effect predictors. This outcome was particularly prominent in the GLMM examining the overall association between vector species richness and malaria prevalence where random effects accounted for more than 30% of the variation in malaria prevalence across observations (amount of variance in a response variable explained by random effects is given by R^2^c-R^2^m; Table 2), representing 97-100% of the total variance explained (proportion of total variance explained attributable to random effects is given by 1-(R^2^m/R^2^c); Table 2). This is perhaps unsurprising when working with data spanning a large spatial extent and long time period pooled together. One notable exception was for vector species richness in sub-Saharan Africa, where all spatio-temporal variation in vector richness at the scales measured was captured in how the environment modified vector communities (R^2^c=R^2^m; Table 3), contrasted with malaria prevalence where much of the variation was explained by random effects instead (amount of variance explained by random effect is 30.43%, representing 68.09% of total variance explained; Table 3). For Southeast Asia, however, we see a slightly different pattern, where there was important spatio-temporal variation in vector communities and, to a lesser extent, in the prevalence of malaria, that was not captured by fixed effect predictors (87.96% and 79.49% of total variance in vector species richness and malaria prevalence was explained by random effects, respectively; Table 3). A more detailed exploration of the ecology of vector communities at local scales would therefore be an asset to resolving some of the drivers of malaria at these larger scales.

## Discussion

Recent years have seen great progress toward understanding processes that shape infectious disease patterns around the world (Stephens et al. 2016; Murray et al. 2018). Mammalian and avian diversity have been the focus of many such studies, as these taxa commonly serve as alternative hosts to human parasites. For vector-borne diseases such as malaria, vector diversity, which was expected to conform to the same latitudinal gradient of diversity observed for most terrestrial fauna, has long been hypothesized to influence disease distribution in the same manner as their vertebrate host counterparts (Nunn et al. 2005; Clark et al. 2014). Yet, these associations have not been tested and global vector diversity has not been fully mapped. While tremendous efforts have been made to map and predict the presence of individual vector species based on climate and habitat suitability (e.g., Hay et al., 2010), now is the time to extend these approaches to consider entire vector communities and how they function across a range of environments (Park et al. 2015). The harmonization of disease ecology and epidemiology is long overdue (Galvani 2003).

In this study, we showed that vector diversity played a key role in structuring global malaria distribution and found these relationships to be highly context dependent. We uncovered vector distribution patterns that were not consistent with the assumed latitudinal gradient of diversity (Fig. 2B) and not aligned with that for the family *Culicidae*, nor for *Anopheles*, the only genus of mosquito with species that act as malaria vectors (Foley et al. 2007). We demonstrate that the global vector species richness and malaria relationship (negative trend that was not statistically significant; Fig. 4A) is not representative of local trends (positive association weakened with increasing latitude; Fig. 4B) – a case of Simpson’s Paradox (Allison and Goldberg 2002). Our SEM analysis further revealed striking differences in environment-diversity-disease relationships emerging from distinct transmission contexts across two malaria endemic regions, sub-Saharan Africa (Table 3, Fig. 5A) and Southeast Asia (Table 3, Fig. 5B). These insights present an opportunity to chart a new path forward in vector-borne disease research where patterns captured at large spatial scales may be used to motivate local investigations and control efforts, as well as guide research in emerging and understudied vector-borne diseases (Murray et al. 2015).

The decreasing species richness of malaria vectors from the edge of the tropics towards the equator (per site-level records; Fig. 2B) is unexpected given the large number of *Anopheles* species apparently found near the equator (per country-level records; Foley et al., 2007). This pattern may arise if low latitude areas were under-represented in our dataset, but that is not the case (Fig. 2), and it also seemed unlikely that malaria vectors would be systematically under-sampled compared to non-vector Anophelines. This poses the intriguing question of why those Anophelines near the equator are not considered vectors. If there are in fact reservoirs of *Anopheles* mosquitos of unknown vector competency residing outside of human settlements where surveys are typically carried out, then this should be a cause for concern (Epopa et al. 2020; Cross et al. 2021). Many human activities and interventions, such as land development and vector management programs, have the potential to alter vector community structure. These changes are often idiosyncratic and depend on exactly how the habitat was altered, as well as which species were available for recolonization in the broader regional pool (Yasuoka and Levins 2007). Unfortunately, publicly available malaria vector survey data often have sparse spatial and temporal coverage (Fig. 2 data points span 1990-2005) and rarely include secondary vectors. In this current collection, we extended the Malaria Atlas Project (MAP) database by adding in observations of secondary vectors when these were present in the original studies compiled in MAP. However, as MAP by design omits secondary vectors from its records, we do not have any information on surveys where only secondary vectors were found, and most country-level records (as used in Foley et al. 2007) are compiled at spatial and temporal scales that cannot be compared to MAP data. The paucity of knowledge of the spatial ecology of most malaria vector species, let alone those thought to be of lesser importance, severely hinders the ability to accurately predict and respond to evolving malaria risks (Fernandes et al. 2018; Russell et al. 2020). More funding and human resources should be devoted to surveying vector communities, in particular, to establish long-term monitoring programs and to extend sampling to non-conventional locations. Results should be properly archived and freely disseminated whenever possible so that researchers may collectively extract more knowledge and maximize returns on these hard-earned datasets.

The observation of few competent vectors at low latitudes led to the inference of a negative association between vector species richness and malaria prevalence at the global scale (Fig. 4A). When this relationship was broken down by latitude, however, we found that vector species richness was in fact positively associated with malaria prevalence near the equator, and the effect weakened and reversed moving toward the edge of the tropics (Table 2, Fig. 4B). These patterns remain quantitatively similar regardless of whether secondary vectors were included in the model (Table 2), suggesting that they were not driven by the occurrence of less-competent species at higher latitude sites. Rather, this is a clear signal of context-dependency in vector diversity and disease relationships: variation in environmental conditions may support vector communities of different sizes and composition (e.g., Parham & Michael, 2010; Fig. 1A, top), or may modify their disease transmission capacities independent of any effects on vector community structure (Park et al., 2015; Fig. 1A, bottom). That the strength and nature of biological interactions can vary depending on the latitude has been observed in a wide range of ecological systems, including hosts and their parasites (Schemske et al. 2009). These principles serve as an essential pillar of disease macroecology theory and are important to consider in future studies.

The two major malaria endemic regions in the world, sub-Saharan Africa and Southeast Asia, are known to differ broadly in their malaria epidemiology, vector community structure, and underlying disease transmission contexts (Baird, 2017; Carter & Mendis, 2002; Table 1; Fig. A1 in Appendix A). Of particular relevance to this study were the differences in vector assemblages: their membership did not overlap, with sub-Saharan African vector communities dominated by a few human-specialist species, and Southeast Asian vectors communities tending to be made up of generalist and opportunistic vectors (Sinka et al., 2010, 2011; Table 1, Fig. 3). Although vector species richness was positively associated with malaria prevalence in both regions, vector community composition only had an effect in sub-Saharan Africa but not Southeast Asia (Table 3, Fig. 5). One explanation of this pattern is that competence among vectors is much more even in Southeast Asia, thus the disease transmission capacity of vector communities would depend on the number of species present but less so on their identities. These community-level characteristics likely also contribute to the observed variation in how vectors functioned in their respective environments (Fig. 5). In sub-Saharan Africa, the environment exhibited a top-down influence on malaria prevalence via vector community structure (Fig. 5A), whereas in Southeast Asia, the environment instead modulated the disease transmission efficacy of vector communities (Fig. 5B). Considering that human specialist vectors mostly blood-feed and rest indoors, it follows that the environment would have little effect on how they perform, but could influence whether a particular species is found. In contrast, in Southeast Asia where vector communities consist mostly of generalists that blood-feed outdoors, their interactions with human hosts are much more likely to be modified by the ambient environment (Stresman 2010). A solid understanding of vector behavioral ecology and a precise definition of vector competence would be essential to further dissect these associations. Here, we employed broad characteristics of vector community structure and distinguished vectors by their primary vs. secondary vector designations, but ideally, we would have a trait-based definition of vector competence and be able to connect these traits with those that determine phenology and distribution. Such data will permit more accurate predictions of how vector-borne disease risk will shift when climate and habitat inevitably change.

Malaria is often considered “a disease of poverty”, where socio-economic disadvantages are consistently linked to increased susceptibility and exposure, and poor disease outcome (Gallup and Sachs 2001; Worrall et al. 2005; Tusting et al. 2013). In this study, we found no direct effect of regional GDP on malaria prevalence (Table 3, Fig. 5), however, we did observe a strong interaction between GDP and vector species richness in Southeast Asia (Table 3, Fig. 5B, Fig. B2-iii in Appendix B). This suggest that high regional GDP can alleviate the positive association between vector species richness and malaria prevalence (Fig. B2-iii in Appendix B), perhaps by modifying the rate of exposure to those vectors (e.g., through better housing structure; Tusting et al., 2015). Despite this strong effect, there was still a large amount of spatio-temporal variance in malaria prevalence that was not captured by this predictor (Table 3, Fig. B2-iii in Appendix B).

Combined with the lack of effect of regional GDP in sub-Saharan Africa, this suggests that finer scale measures of socio-economic condition would be helpful. Regional GDP was conceived here as a proxy of investment into health care and other infrastructure, but in practice is an indicator of economic productivity and may not necessarily be tightly associated with our target phenomenon. Future studies should refine this proxy and explore the many other facets of social-economic development that may interact with climate, habitat change, host movement, and vector community structure to influence disease risk and outcome (Lambin et al. 2010; Kilpatrick and Randolph 2012; Gottdenker et al. 2014; Parham and Hughes 2015). For instance, since wealth at the regional level may not trickle down evenly, it could be useful to consider finer-scale metrics such as household income and housing quality (Tusting et al. 2013, 2015). It would also be valuable to extend the analysis to investigate the influence of social and economic factors on malaria disease outcome, by modelling death rate and disability-adjusted life years as the response variable. The SEM approach adopted here is particularly apt at untangling webs of relationships to reveal critical paths and parameters (Grace et al. 2010; Shipley 2016) – we have but scratched the surface of possibilities with our model (Fig. 1).

Worldwide malaria deaths have declined greatly in the past two decades, a global health triumph attributed in large part to effective vector management (Bhatt et al. 2015; Gething et al. 2016). However, progress towards eradication appears to have stalled in recent years (Alonso and Noor 2017). Current major obstacles include donor fatigue resulting in momentum loss in vector management programs (Nkumama et al. 2017; Russell et al. 2020), and spread of artemisinin resistance in the Greater Mekong Subregion undermining effectiveness of drug treatment (Wongsrichanalai and Sibley 2013; Christofferson et al. 2020). The recent invasion of *Anopheles stephensi* in Ethiopia is also a concern, as it is one of the few urban-adapted malaria vectors and thus could establish in large cities in sub-Saharan Africa (Sinka et al., 2020; Takken & Lindsay, 2019). The future of malaria is being shaped by rapid environmental and societal change. An interdisciplinary systems approach is crucial to continuing to make progress in the race to elimination.

## Materials and methods

### Data sources and management

We curated a dataset of global malaria prevalence and vector distribution by integrating information from several open-access databases. Malaria was defined as infections caused by *Plasmodium falciparum, P. vivax, P. malariae, P. ovale*, and *P. knowlesi*, or combinations of these parasites. Vectors were categorized as being primary (or “dominant”) species, which contribute significantly to the transmission of malaria (Hay and Snow 2006; Service 2012), or secondary species that play a smaller role (Antonio-Nkondjio et al. 2006). While there is a defined list of primary vectors (Hay and Snow 2006; Service 2012), designation of secondary vectors is less straightforward and typically *ad hoc*. Thus, it was not possible to create an exhaustive list of secondary vectors of malaria. Further, much of Anopheline taxonomy is still under debate (Harbach and Kitching 2016) and some species complexes are rarely resolved into distinct species due to phylogenetic uncertainty or technical constraints in the field. The vector species included in this study thus likely underestimate the true diversity of malaria vectors. In spite of these limitations, this dataset contributes to a growing collection of biodiversity-disease databases, which tend to focus on host diversity (e.g., Dunn et al., 2010; Stephens et al., 2017; Wood et al., 2017).

Vector occurrence and malaria prevalence data were extracted from the Malaria Atlas Project (MAP; Hay and Snow 2006; Guerra et al. 2007; Hay et al. 2010). MAP is an open-access database created and maintained by an international consortium of malaria experts. The entomological (primary vectors only) and epidemiological (*P. falciparum* only) surveys extracted from MAP were matched in space (within 1×1 km^2^ of each other) and time (overlapped in study duration) and validated against original sources. Longitudinal data were treated as repeated measures in our statistical analyses when temporally-resolved data were available, however, in most cases longitudinal data were presented in aggregate form (95 records, of which 71 spanned less than a year, and the remainder up to three years), or had to be aggregated to facilitate matching (22 records). Information on vector abundance, observations of secondary vectors, and other *Plasmodium* species were added when available from the primary source. Entries that could not be validated (e.g., personal communications) were removed. Some records (n=5) were retrieved from comparative studies that assessed the efficacy of various malaria control interventions (e.g., bed net trials; Etang et al. 2007) and we retained only data from control sites (or pre-treatment values) from such studies. The resulting matched and verified mosquito community and malaria prevalence data consisted of 163 unique observations from 112 locations.

Environmental variables and a measure of economic condition were added to the dataset for the investigation of factors that mediate vector diversity and disease relationships. Monthly temperature and precipitation at 15 arc-min spatial resolution (approximately 30 km^2^ at the equator) was extracted from Copernicus Earth Observation Programme of the European Union, Climate Change Service (http://climate.copernicus.eu), and matched to the location and month of each vector and disease observation. For aggregated longitudinal studies, the average monthly temperature and precipitation over the study duration was calculated. We then obtained sub-national (i.e., administrative units within a country) per capita gross domestic product (GDP; Kummu et al., 2018) as our measure of economic condition, to serve as a proxy for investment into health services and infrastructure. GDP may directly influence disease prevalence through healthcare expenditure, as well as have disparate effects on mosquito species richness and community composition as vector management programs typically only target primary vectors (Killeen et al. 2017; Waite et al. 2017). This data was provided at 5 arc-min spatial resolution (approximately 9 km^2^ at the equator) for the years 1990-2005. Epidemiological and entomological data that fell outside of this time span were removed (1983-1989, n=30), reducing the dataset to 133 observations from 92 locations.

### Statistical analysis

We first explored global mosquito diversity and malaria prevalence by mapping these attributes (n=133), and analyzed the representation of primary vector species relative to all vectors based on total species richness and abundance with a subset of the site-level data where species abundances were available (n=83). We then used generalized linear mixed models (GLMM) to test for associations between vector species richness and malaria prevalence. Only sites located within the tropics were included in this (and all subsequent) analysis as most observations (126 out of 133) were made there. The response variable, malaria prevalence, was modelled with a binomial distribution weighted by the number of individuals sampled (see Appendix C for a note on model fit diagnostics). The focal predictors were the number of vector species observed and the interaction between latitude and vector species richness. In total, we constructed four models that included the effects of primary or all vector species richness, and with or without latitude by species richness interaction, in a factorial manner. This setup allowed us to investigate the following two questions: 1) is the effect of vector species richness predominantly driven by primary vectors, or does inclusion of secondary vectors better explain malaria prevalence (by contrasting models with primary vs. total vector species richness), and 2) is the effect of vector species richness general, or does its effect depend on the environment (by contrasting models with and without latitude by vector species richness interaction)? We used the absolute value of latitude (i.e., distance from equator) because changes in environmental conditions from the equator to the poles are expected to be roughly symmetrical about the equator (Hillebrand 2004). All predictor variables were scaled to unit variance to facilitate model convergence. Models also included study site, country, and year, nested in succession, as random effects to account for the hierarchical structure of the data.

To specifically dissect the environmental factors that mediate vector diversity and disease relationships, we developed structural equation models (SEMs) that explicitly account for the potential associations between factors that may otherwise pose issues of multicollinearity in simple regressions. We included two measures of vector diversity in our SEM: total number of vector species observed and percentage that were primary vectors (a proxy for community composition). These attributes were correlated with each other, but each may independently influence malaria prevalence – SEMs allow us to distinguish between them. We included monthly temperature and rainfall as environmental predictors, as well as GDP as an economic predictor, of vector species richness, percent primary vectors, and malaria prevalence. Lastly, we included interactions between abiotic and biotic factors in a factorial manner (six interaction terms total). The structure of the full SEM is depicted in Fig 1B.

We evaluated this SEM with the R package *piecewiseSEM* (Lefcheck 2016). This approach deconstructs SEMs into a collection of regression models, one for each response variable and factors hypothesized to influence it, and therefore allows for non-normal distributions and hierarchical structure to be incorporated into each submodel. There are three factors that act as responses to other variables in our SEM: malaria prevalence, vector community composition, and total vector species richness. These responses were modelled with binomial, gaussian, and Poisson distributions, respectively (see Appendix D for a note on model fit diagnostics). All submodels have the same hierarchical structure as the previously described GLMMs.

Due to inherent difficulties of fitting a complex SEM to a relatively small dataset, we introduced model selection and reduction procedures prior to evaluation (after Roland et al., 2019). We first optimized each submodel by fitting a series of models that represent all possible combinations of predictors from the full set (i.e., we “dredged” each submodel). Predictors from models with AIC scores within two units of the lowest value were retained for subsequent analyses. A submodel was removed from the SEM when the intercept-only model was the “best” model (i.e., the intercept-only model has a lower AIC score than the next best model by 2 units or more), and when its associated response variable did not predict another factor. The “best” submodels were then incorporated into an SEM and subjected to the test of directed separation (Shipley 2000) to ensure no important paths were missing. We used this procedure to identify the “best” model for sites in Africa and Asia separately, resulting in distinct SEMs for these regions.

## Acknowledgements

We thank past and current members of the Mideo lab – Dr. Max Farrell, Dr. Megan Greischar, Dr. Cylita Guy, Madeline Peters, Dr. Tsukushi Kamiya, Antonio Lorenzo, Dr. Alexander Whitlock – for valuable discussion and moral support throughout this project. We are also grateful to Dr. Marla Sokolowski for feedback on this work.

