## Appendix A-D for "Vector diversity and malaria prevalence: global trends and local determinants"

Here we provide more information on the dataset we collected. Figure A1 presents a comparison of site characteristics across the two study regions, sub-Saharan Africa and Southeast Asia. These factors varied across the regions in a manner consistent with the literature (see Table 1).

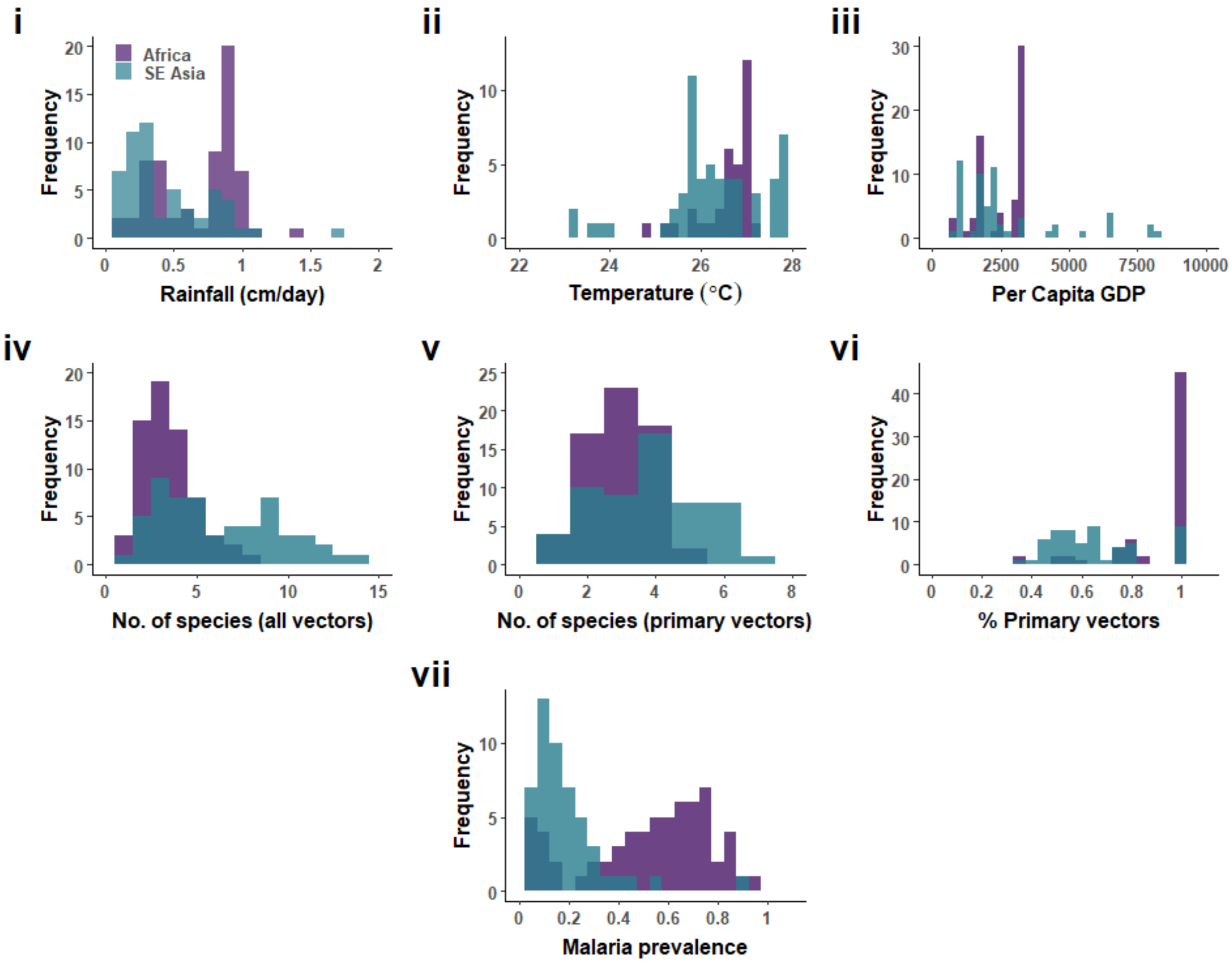


**Fig A1. Frequency distributions of i) rainfall, ii) temperature, iii) per capita GDP in 2010 USD, iv) total vector species richness, v) primary vector species richness, vi) percentage of species that were primary vectors, and vii) malaria prevalence across site in sub-Saharan Africa (n=64) and Southeast Asia (n=57).**

### Appendix B

To aid with the interpretation of the structural equation model results (Table 3 and Fig. 5 in main text), each statistically significant association was graphed independently below. Fig. B1 depicts the influence of environment (rainfall) on vector species richness in African sites, which in turn influenced malaria prevalence. The proportion of species that are primary vectors also had a positive effect on malaria prevalence. Fig. B2 depicts the effects of the interactions between abiotic (rainfall and GDP) and biotic (vector species richness and percent primary vectors) factors on the prevalence of malaria in Southeast Asia. In Fig B2, panels (ii) and (iii), the discrepancy between the raw data points and the model-estimated relationships reflect the substantial spatial-temporal variation in malaria prevalence not captured by the fixed effects in these models.

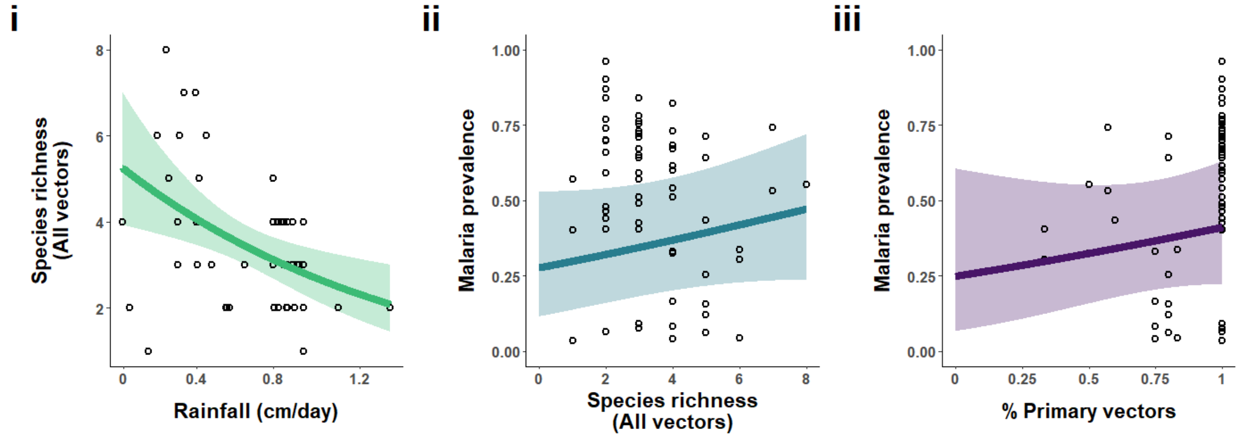


**Fig B1. i) Predicted effect of rainfall on vector species richness in African sites (n=64).** Vector species richness was modelled using generalized linear mixed models with Gaussian distribution. **ii-iii) Predicted effects of vector community attributes on malaria prevalence (n=64).** Malaria prevalence was modelled using generalized linear mixed models with binomial distribution weighted by the number of individuals sampled. All models include study site, country, year, nested in succession, as random effects to account for the hierarchical structure of the data. Bands represent 95% confidence interval of prevalence estimates, and points represent original data.

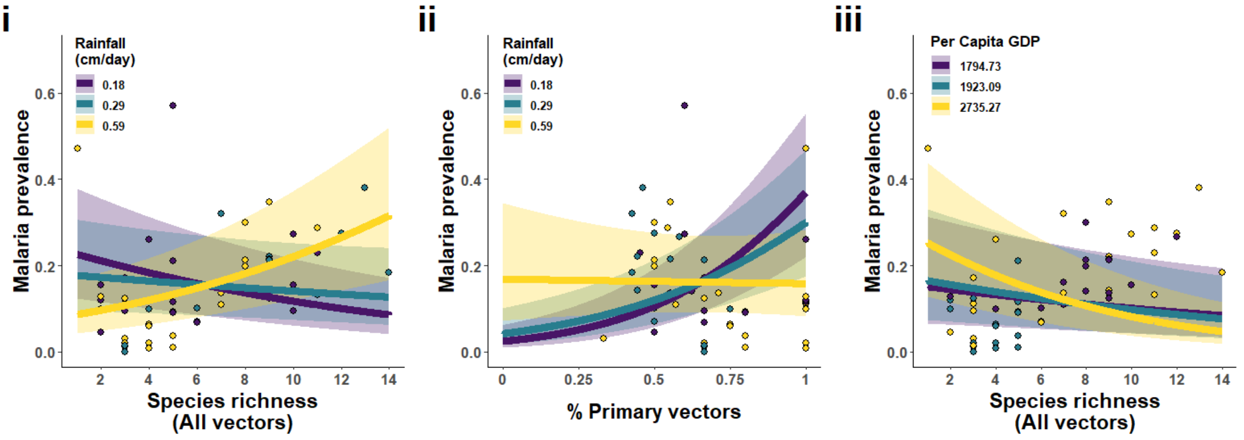


**Fig B2. Predicted effects of various vector community attributes and abiotic interactions on malaria prevalence in Southeast Asian sites (n=57).** Malaria prevalence was modelled using generalized linear mixed models with binomial distribution weighted by the number of individuals sampled. Predictor variables considered were **i) the interaction between vector species richness and rainfall, ii) the interaction between proportion of primary vectors and rainfall**, and **iii) the interaction between vector species richness and per capita GDP**. All models include study site, country, and year, nested in succession, as random effects to account for the hierarchical structure of the data. Within each panel, the values chosen for the environmental variable depicted (rainfall and GDP) correspond to the 1$^{st}$ quartile (purple), median (teal), and 3$^{rd}$ (yellow) quartile of the range of values observed at the Southeast Asian sites (Appendix A, Fig. A1). Bands represent 95% confidence interval of prevalence estimates, and points represent original data cut into three equal-length bins (i.e., each category consist of the same number of data points, and corresponds to the first, middle, and third quartiles of the range of values).

### Appendix C

In this appendix, we present and discuss the steps taken to validate the model assumptions of the GLMMs used to assess the global association between malaria prevalence, vector species richness, and latitude (Table 2 in main text). Figure C1 shows the Quantile-Quantile (Q-Q) plot used to assess the assumption of homogeneity of variance in two models that included the interactions between primary vector species richness and distance from equator (absolute latitude; panel i) and between total vector species richness and distance from equator (panel ii). Both Q-Q plots indicate there was some violation of this assumption, and the pattern shown suggested that it was due to overdispersion in our data, which could inflate our Type I error rate (i.e., false positive rate). We therefore adjusted our results for overdispersion with the procedure outlined in Faraway (2005), which involved adjusting the standard error as a function of the dispersion factor and recomputing the p-value. We found that results are qualitatively unchanged following this adjustment (Table C1). We are therefore confident in our conclusion that the interaction between total vector species richness and distance from equator is a statistically significant predictor of malaria prevalence (marginally significant in the case of primary vector species richness).

**Table C1. Generalized linear mixed models for malaria prevalence at 126 sites across the tropics, corrected for overdispersion.** The response variable for all models was malaria prevalence, coded as an aggregate binomial variable and weighted by number of individuals sampled. The following predictor variables were considered: vector species richness (primary vectors only or all vectors), distance from the centroid of each site to the equator (i.e., absolute value of latitude), and their interaction. All predictors were centered and scaled to unit variance to facilitate model convergence. Several random effects were included to account for non-independence in the data: site (some sites were measured repeatedly), country, and year, nested in that order. β: regression coefficients. SE: standard error. Model output metrics, including standard errors for effect size estimates and p-value, were adjusted for overdispersion based on procedure outlines in Faraway (2005).

| | **Primary vectors only** | | | **All vectors** | | |
|---|---|---|---|---|---|---|
| **Predictors** | **β** | **SE** | **p-value** | **β** | **SE** | **p-value** |
| Intercept | -1.03 | 0.40 | 0.010 | -1.046 | 0.40 | 0.009 |
| Species richness | 0.14 | 0.09 | 0.11 | 0.19 | 0.16 | 0.23 |
| \|Latitude\| | -0.23 | 0.28 | 0.42 | -0.23 | 0.29 | 0.44 |
| Species richness * \|latitude\| | -0.16 | 0.08 | 0.032 | -0.26 | 0.13 | 0.052 |

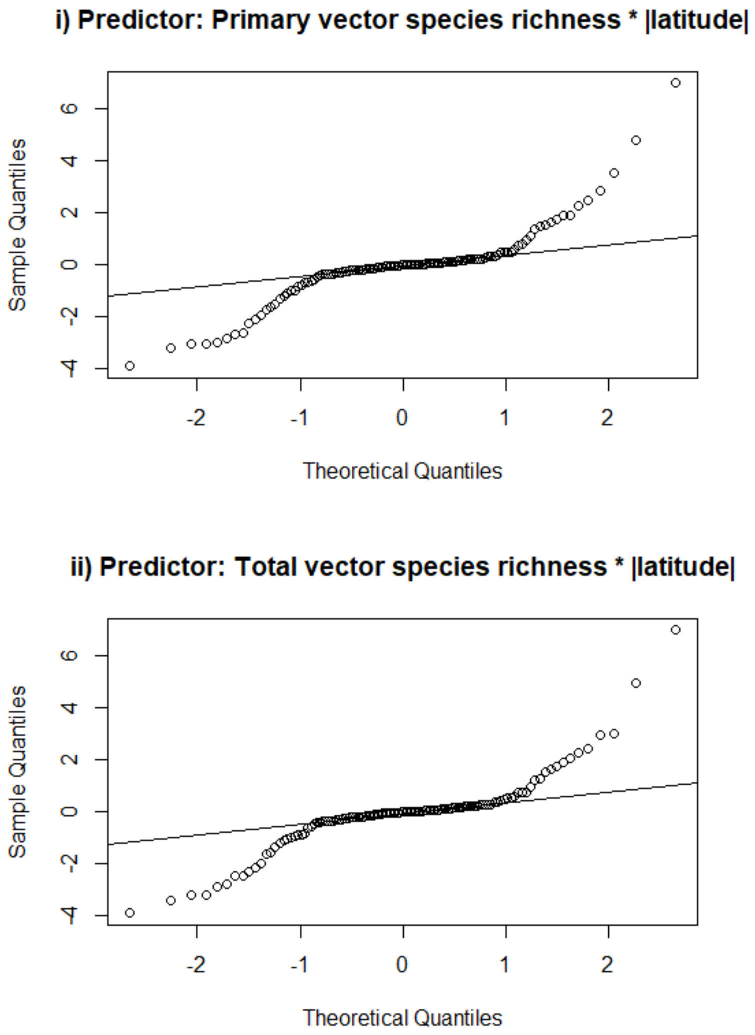


**Fig C1. Quantile-quantile (Q-Q) plots for diagnosing GLMM assumptions.** The response variable was malaria prevalence in both models. For the homogeneity of variance assumption to be valid, all data points should appear close to the solid line. Deviations from the solid line at the tails, as seen above, typically suggest the data are overdispersed.

**Appendix D**

We used structural equation models (SEM) to dissect the various pathways through which environmental and vector community attributes influenced each other and, in turn, malaria prevalence (Figure 1 in main text). There were three sub-models embedded in this SEM. One of these models had vector community composition (percent of vectors species that are primary vectors as a proxy) as a response variable, modelled with a Gaussian distribution. We inspected the validity of the assumption of homogeneity of variance for this model and found that there was some violation (Figure D1-i). However, refitting the model with a binomial distribution (which is the more common approach for this type of data) did not improve model fit, and in fact, appeared to result in greater deviations from the assumption (Figure D2-ii). Transforming the response variable also did not improve model fit (results not shown). We note, however, that the alternative model structure did give different results: in the Gaussian model, vector community composition was not significantly associated with any of the predictors considered, but in the binomial model temperature and rainfall became statistically significant predictors of vector community composition (Table D1, Figure D2). Despite this difference, the overall inference of the SEM – in sub-Saharan Africa, the environment influences malaria prevalence indirectly via its effects on vector community structure – remains the same across the two models (Figure D2 compared with Figure 5A in main text). We therefore chose to present the SEM where vector community structure was modelled with a simpler structure (Gaussian distribution) because alternative model structures did not improve model fit, and did not result in qualitatively different inferences from the SEM.

**Table D1. Results for structural equation model fitted to sites in sub-Saharan Africa following model selection.** All response variables, along with all corresponding predictors as depicted in Figure 1B (main text) are listed below. All sub-models also include a random effect term to account for non-independence in the data: site (some sites were measured repeatedly), country, and year, nested in that order. This model differed from the ones presented in Table 3 (main text) in that vector composition (percent of vectors that are primary vectors as proxy) was modelled with a binomial distribution (number of primary vector species weighted by total number of species observed) instead of Gaussian. A blank cell indicates a path that was dropped during model selection (via procedure detailed in main text). Raw effect estimates (Raw β), standardized effect estimates (Std β), and standard errors are provided. The d-separation test was used to assess model fit, and the associated Fisher's C statistic, degrees of freedom, and p-value are reported. The null hypothesis of the d-separation test is that there are no important links missing in this model. Coefficients of determination ($R^2$) for the focal response variable (malaria prevalence) are given. Conditional $R^2$ ($R^2c$): proportion of variation explained by fixed and random effects combined. Marginal $R^2$ ($R^2m$): proportion of variation explained by fixed effects only. $R^2$ statistics were estimated using methods outlined in Nakagawa and Schielzeth 2013, and Nakagawa et al. (2017).

| Response | Predictor | Raw β | Std β | SE |
|---|---|---|---|---|
| Malaria prevalence | Species richness | 0.20† | 0.14† | 0.09 |
| | Composition | 3.65† | 0.28† | 1.52 |
| | Rainfall | | | |
| | Temperature | 0.030 | 0.013 | 1.00 |
| | GDP | | | |
| | Richness * rain | | | |
| | Richness * temp | -0.08 | -0.14 | 0.09 |
| | Richness * GDP | | | |
| | Composition * rain | | | |
| | Composition * temp | 0.89 | 0.33 | 0.89 |
| | Composition * GDP | | | |
| Species richness | Rainfall | -0.32† | -0.74† | 0.12 |
| | Temperature | 0.08 | 0.20 | 0.10 |
| | GDP | 0.47 | 0.18 | 0.62 |
| Composition | Rainfall | 0.83† | 0.37† | 0.34 |
| | Temperature | 0.56† | 0.25† | 0.26 |
| | GDP | | | |
| **Correlations** | | | | |
| Species richness | Composition | | -0.56† | |
| Temperature | Rainfall | | 0.72† | |
| GDP | Temperature | | 0.62† | |
| GDP | Rainfall | | 0.71† | |
| **Goodness of fit** | | | | |
| Fisher's C | | | 2.35 | |
| Degrees of freedom | | | 6 | |
| p-value | | | 0.88 | |
| Malaria $R^2c$ | | | 40.14% | |
| Malaria $R^2m$ | | | 9.71% | |
| Species richness $R^2c$ | | | 16.99% | |
| Species richness $R^2m$ | | | 16.99% | |
| Composition $R^2c$ | | | 22.06% | |
| Composition $R^2m$ | | | 22.06% | |

†p<0.05

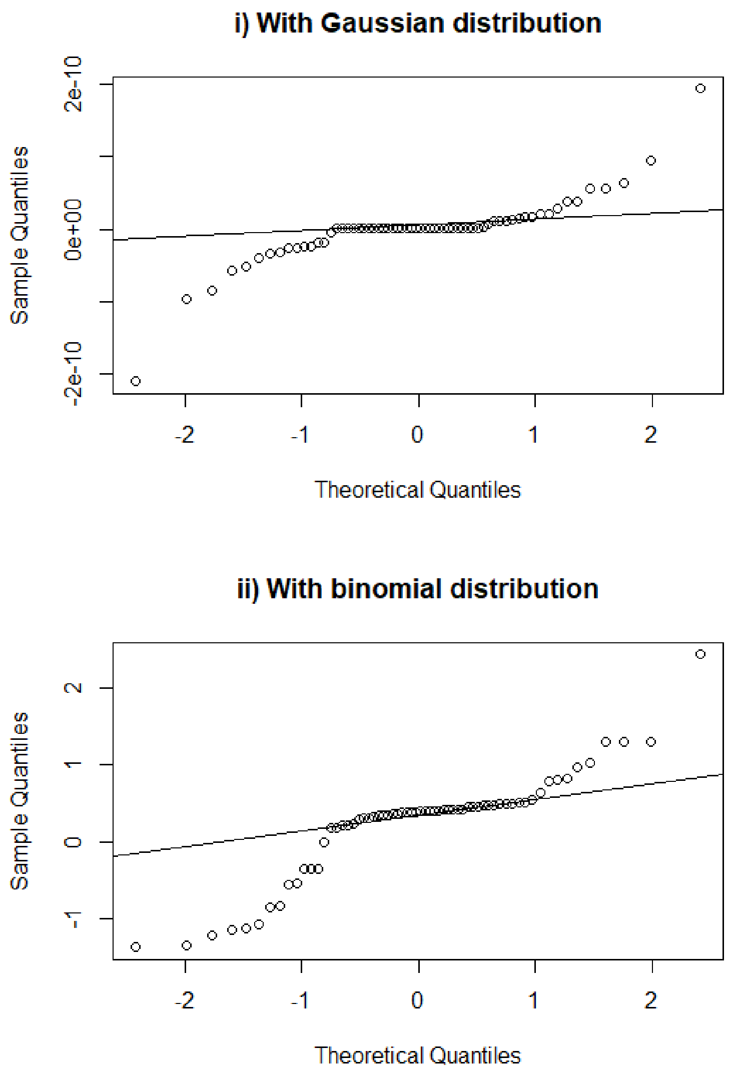


**Fig D1. Quantile-quantile (Q-Q) plots for diagnosing SEM sub-model assumptions.** For both models, the response variable was percentage of vector species that are primary vectors, and predictor variables were rainfall, temperature, and regional GDP, to facilitate comparison of the different distributions used. For the homogeneity of variance assumption to be valid, all data points should appear close to the solid line. The above Q-Q plots show neither model meets this assumption, as the data points stray from the theoretical line at the ends.

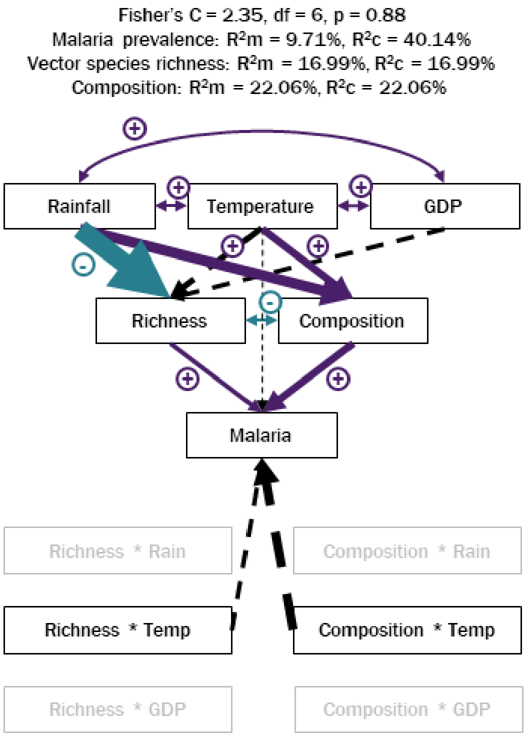


**Fig D2. Path diagrams depicting model selection and fit results for sub-Saharan Africa data subset (n=64).** Model selection procedure detailed in the main text. This model differed from the ones presented in Figure 5A (main text) in that vector composition (percent of vectors that are primary vectors as proxy) was modelled with a binomial distribution (number of primary vector species weighted by total number of species observed) instead of Gaussian. Paths (i.e., arrows) represent the direction of the modelled causal relationship between variables, with the arrow pointing from the predictor to the response. Arrow size is proportional to the strength of paths. Double-headed arrows indicate correlations and are not scaled to effect size. Purple arrows represent positive associations, teal represent negative associations, and dashed arrows are associations that were not statistically significant. Variables in grey were dropped during model trimming. See Table D1 for details of model fit results. Fisher's C statistic and associated p-value from model goodness-of-fit tests, and marginal and conditional $R^2$ ($R^2$m and $R^2$c, respectively) of the response variables provided within the figure.
